# Modulation of optical speaker using biogenic guanine platelets floating in water

**DOI:** 10.1101/2021.09.16.460721

**Authors:** Masakazu Iwasaka

**Affiliations:** Hiroshima University, 1-4-2 Kagamiyama, Higashihiroshima, Hiroshima 739-8527, Japan

## Abstract

Microphones are miniature devices for sound detection. Various technologies have been developed to transfer sound properties onto other physical quantities, e.g., electricity. Over the past three decades, many studies have reported on optical sensing of sound. Most of these studies were performed via application of light interference at the edges of optical fibers. Several studies have reported on detection of sounds in the air or objects causing mechanical vibrations based on light interference. This work proposes an optical speaker which is a method to reconstruct and modulate sound from the power spectrum of light that has been reflected by guanine platelets floating in water droplet. The water droplet containing fish guanine platelets was placed on a piezoelectric membrane and acoustic vibration from the membrane propagated inside the droplet. A photomultiplier tube (PMT) then collected the light reflected from the water droplet. Without post-analysis of the measured light intensity, the analog output voltage from the PMT clearly sounded an audio speaker. In addition, it was found that the guanine platelets reflecting light operated as an audio equalizer.

## Introduction

Huge number of studies have been undertaken on microphones as a representative technology for audible sound detection.^1)–3)^ As a type of electroacoustic technology, microphones contain a transduction unit that transfers the pressure and frequency of sound onto other physical quantities, e.g., electric charge.^2,3)^

Over the past two decades, use of light to form a microphone has been reported in numerous works.^4–11)^ Most previous studies on optical microphones have focused on fiber optic-type measurement systems.^5–8)^ Among the various types of optical measurement technique that were proposed, light interference was useful for detection of aeroacoustic signals.^9–13)^ One of the techniques developed in this period achieved detection of sounds propagating in the air, far away from the optical sensor, without use of optical fibers by applying Fraunhofer diffraction.^10)^ A similar approach involving measurement of laser light interference generated by incident light and reflected light, was also reported.^10–12)^ Noncontact sound detection based on the principle of the Fabry-Pérot interferometer was also achieved.^12)^ Post analysis of the detected signal from an optical microphone using a deep neural network was found to enhance speech signal acquired by an optical microphone.^13)^

Another successful optical sound detection method involved analysis of captured images obtained using a high-speed camera.^14)^ This method, which was named the ‘visual phone’ method, can extract acoustic vibrations from a targeted object. When detecting sound using the effects of acoustic waves on the spatial distribution of the light intensity, it is possible for the acoustic waves to act as a transducer and transfer the frequency information from an acoustic wave into a light intensity vibration. For example, it was reported that wing of Morpho butterfly acted as a vibrator for acoustic sensing.^15)^

Based on this viewpoint, the method presented in this study is designed to use a water droplet as a sound-to-light-intensity transducer. The surface vibrations of the water droplet containing guanine platelets under audible sound exposure are investigated with the aim of developing an effective sound-to-light-intensity signal transducer. Rather than use laser light, a light-emitting diode (LED) is used as the light source, and a photomultiplier tube (PMT) and a microscope lens are equipped to act as the detection optics. In addition, effects of the addition of biogenic fish guanine platelets to the water droplet was investigated. A technique of bio-photonic audio information carrier transmission on light intensity using guanine platelets was developed.

## Materials and Methods

Figure 1 shows diagrams of the experimental setup used to transfer sound into a light-intensity-vibration *via* a water droplet. Two types of experimental system, i.e., with and without a lens, were used. As shown in Fig. 1(a), a piezoelectric membrane (Thrive piezoelectric micro-speaker unit; resonance frequency of 940 Hz, ϕ35-50u(B)) had the role of generating the audible sound. The membrane surface was painted with black ink to reduce light reflections and maintain the surface hydrophobicity.

**Fig. 1.**
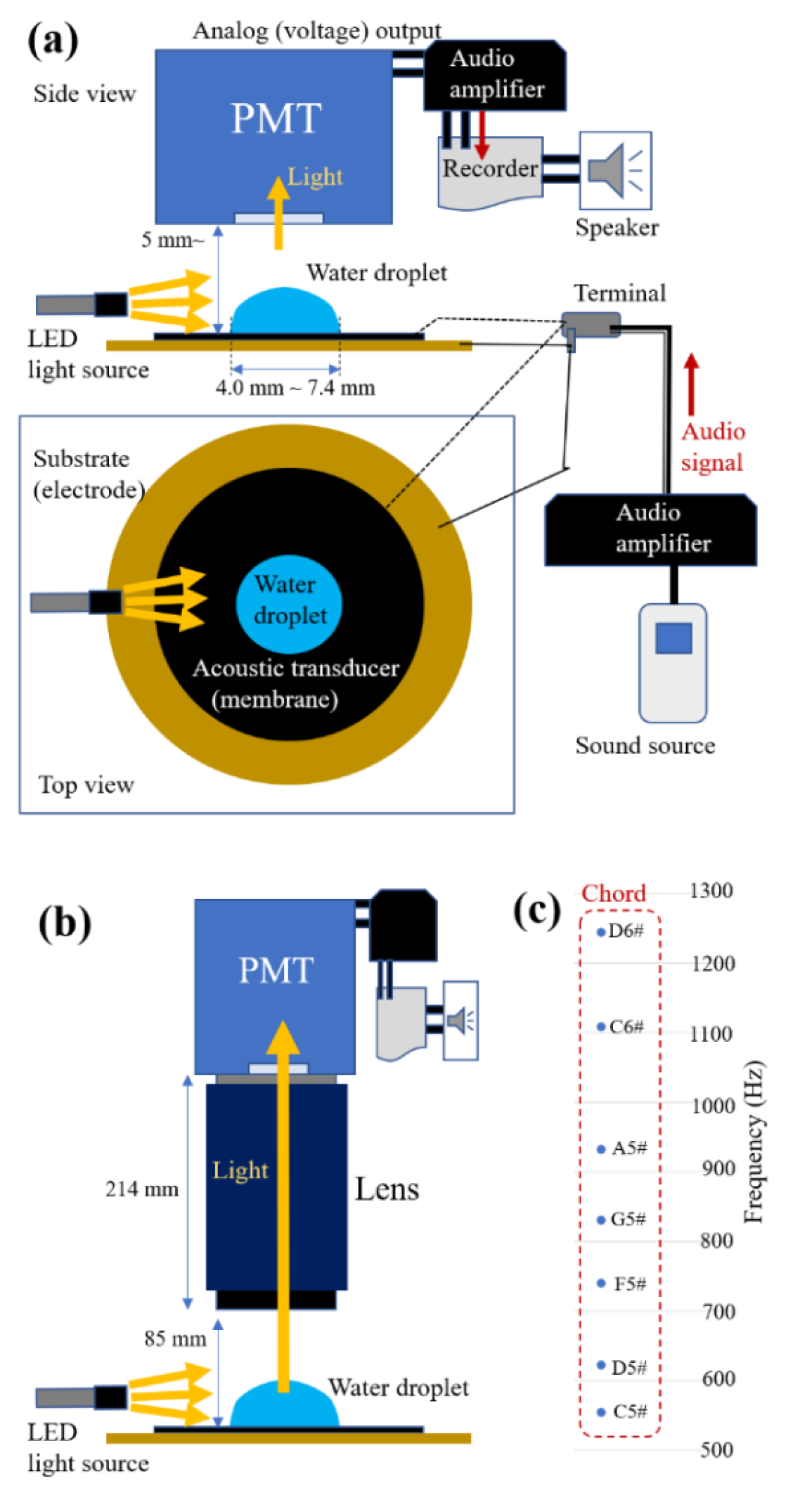
Experimental setup to transfer sound into a light-intensity-vibration *via* a water droplet. (a) Illustration of the system directing light to a PMT-type photon sensor from a water droplet mounted on a piezoelectric membrane (audible sound speaker, i.e., an acoustic transducer). (b) Diagram of the proposed system with a lens. (c) Architecture of the notes in the chord used in the sound source that was applied to the acoustic transducer.

A droplet of distilled water with a volume in the range from 10 μl to 60 μl was placed on the membrane. A white Hayashi Repic LA-HDF158A LED light was used as the light source. The light was passed through a light guide and then reflected by the water droplet, and a PMT (Optomechatoro USB-PMT-S1) collected the reflected light. This PMT was equipped with an analog voltage output circuit that was connected to an audio amplifier (Behringer P2), a pulse code modulation (PCM) recorder (Olympus LS-P4), and a speaker unit.

Fish guanine platelets were separated from scales of goldfish. The suspension of water containing the guanine platelets was utilized as well as the distilled water droplet. The experimental methods used with the fish were approved by Hiroshima University Animal Care and Use Committee (approval numbers F16–2 and F20-4, Hiroshima University).

Figure 1(b) shows a diagram of the modified measurement system with a lens (Navitar 2.0× 1-51473 microscopic lens). This measurement system enabled adjustment of the size of the detection area for the reflected light coming into the detection inlet of the PMT. In addition, use of this lens made it possible to detect sound optically over long distances.

An amplified audio signal was supplied by the audio amplifier (Yamaha PX10 power amplifier) to the two electrodes of the piezoelectric micro-speaker. The sound source amplified by the amplifier was the chord shown in Fig. 1(c), which was pre-recorded using another PCM recorder (Olympus LS-P4). The chord was produced using an electronic musical keyboard (Casio CTK-671 musical keyboard) with its timbre to set to ‘drawbar organ.’ The output voltage supplied to the piezoelectric membrane was in the 8 V to 12 V range, and the membrane sounded using soft tones. The frequency and intensity of the original recorded sound and the optically transferred sound were analyzed using audio spectrum analysis software (Wavosaur 1.7.0.0).

## Results and Discussion

Figure 2(a) shows the power spectrum of the original sound, for which the architecture of the notes in the chord is shown in Fig. 1(c). The horizontal axis (frequency) ranges from 10 Hz to 5000 Hz on a logarithmic scale. Figure 2(b) shows an optically reconstructed spectrum of the sound that passed through the water droplet that was generated from the PMT’s analog voltage output. The sensor head of the PMT was positioned above the water droplet. The distance between the PMT sensor head and the water droplet was approximately 5 mm. From comparison of Fig. 2(a) with (b) it was obvious that the five notes were consistent with the corresponding notes in the original sound. As shown in Fig. 2(a), the timbre of the musical keyboard that was used also produced additional tones that were higher or lower than the seven notes of the chord. These additional tones also appeared in the optically transferred sound (Fig. 2(b)). Figure 2(c) shows the power spectrum of the sound that was transferred from the dry piezoelectric membrane without the water droplet. The spectrum in Fig. 2(c) contained fewer peaks when compared with the other spectra. The recorded audible sound illustrated in Fig. 2(b) showed a clear sound that conformed with the original sound, while the sound illustrated in Fig. 2(c) could not be sensed by healthy human ears.

**Fig. 2.**
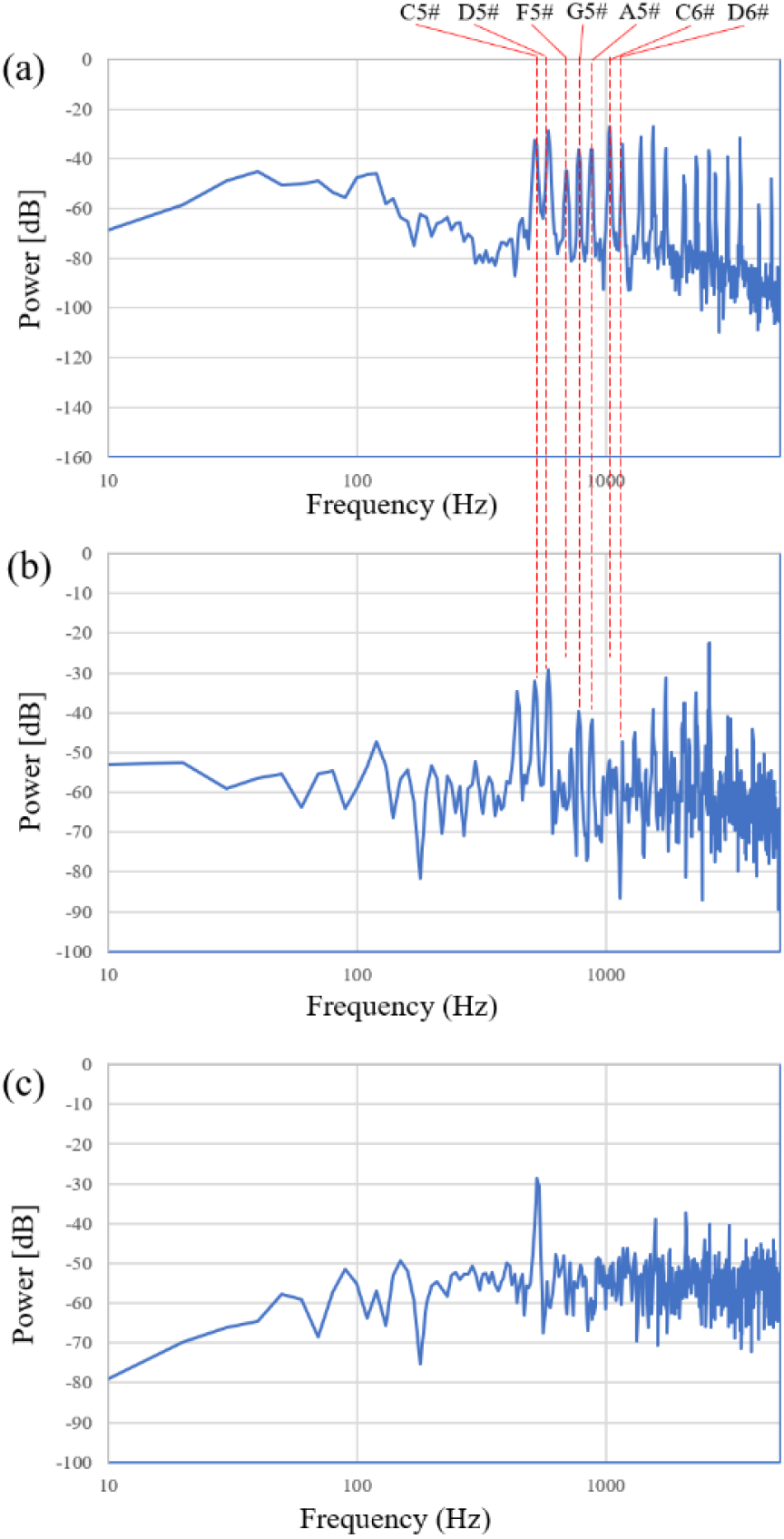
Comparison of the power spectrum of the optically transformed sound transmitted *via* the water droplet with that of the original sound. These measurements were carried out without the lens (i.e., the method shown in Fig. 1(a)). (a) Original sound (*Listen* Supplementary_sound_1). (b) Sound acquired *via* the water droplet (*Listen* Supplementary_sound_2). (c) Sound acquired from the dry piezoelectric membrane without the water droplet.

In Fig. 3, the sound reconstruction was achieved using the PMT when it was equipped with a microscope lens, which was inserted in front of the PMT sensor head. The reconstructed sound was clearer when compared with that reconstructed without the lens. The effect of collecting the reflected light using the lens becomes apparent when the power spectra shown in Fig. 3 are compared with the spectrum shown in Fig. 2(b). The spectra shown in Fig. 3(a)–(d) were obtained in water droplets of different volumes. The water droplets set on the hydrophobic surface formed circular boundaries on the membrane with diameters ranging from 4.0 mm to 7.4 mm. Varying the volume caused the surface inclination and the detected area to be modified. These conditions then affected the audible sound waves that are shown as power spectra in Fig. 3(a)–(d). Each spectrum contained unique peaks as a consequence of this modification of the frequency properties of the sound. Figure 4 shows the dependence of the averaged power on the water droplet volume. The power of the sound within the range from 10 Hz to 5000 Hz increased gradually when the volume of the water droplet increased.

**Fig. 3.**
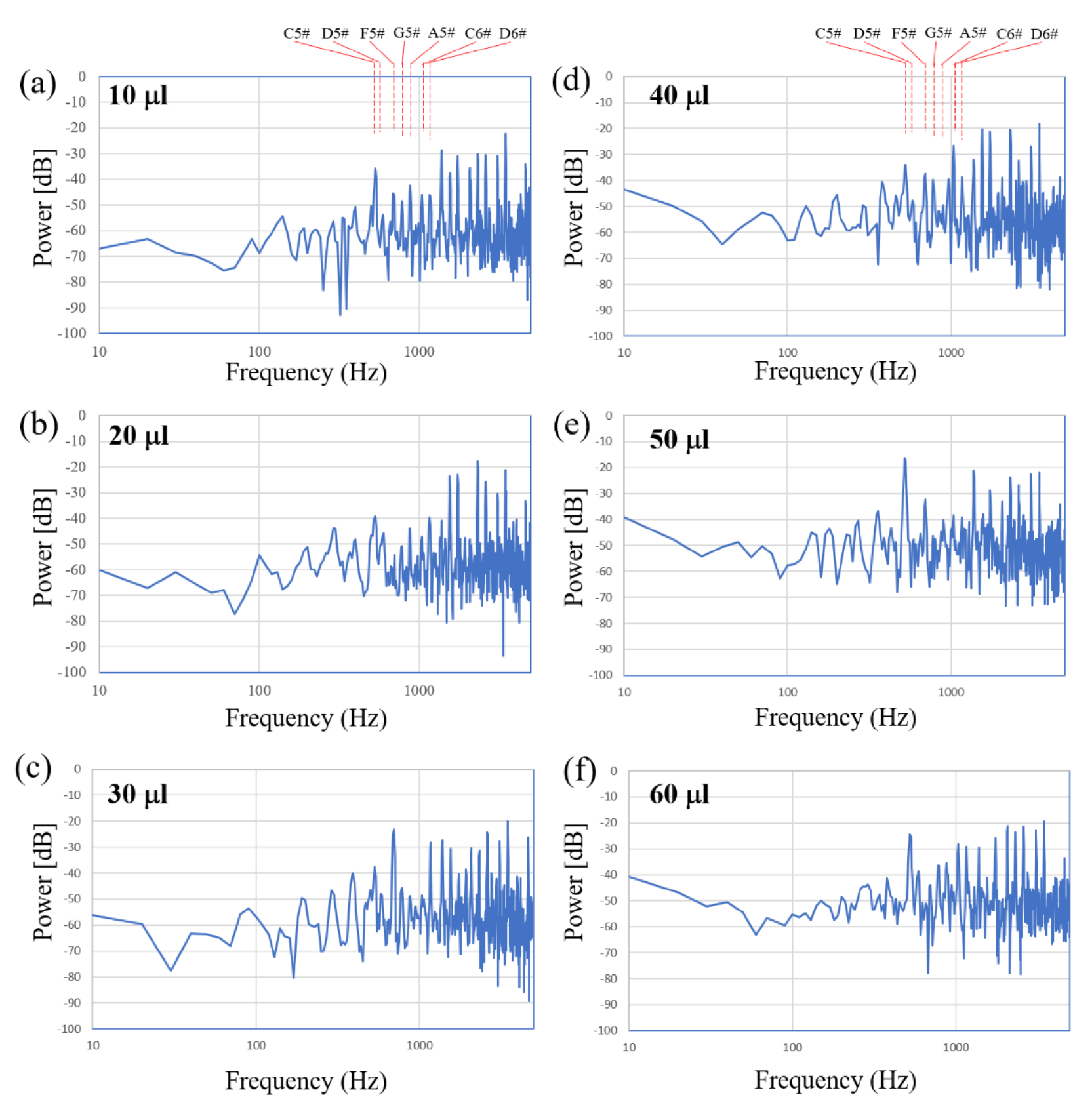
Power spectra of the optically transformed sound acquired *via* the water droplet when detected through a microscope lens. A dependence of the power spectrum profile on the volume of the water droplet was observed. (a) Initial droplet (10 μl). (b)-(f) Volume of the droplet was increased by every 10 μl. The optically transferred sound was continuously recorded (*Listen* Supplementary_sound_3).

**Fig. 4.**
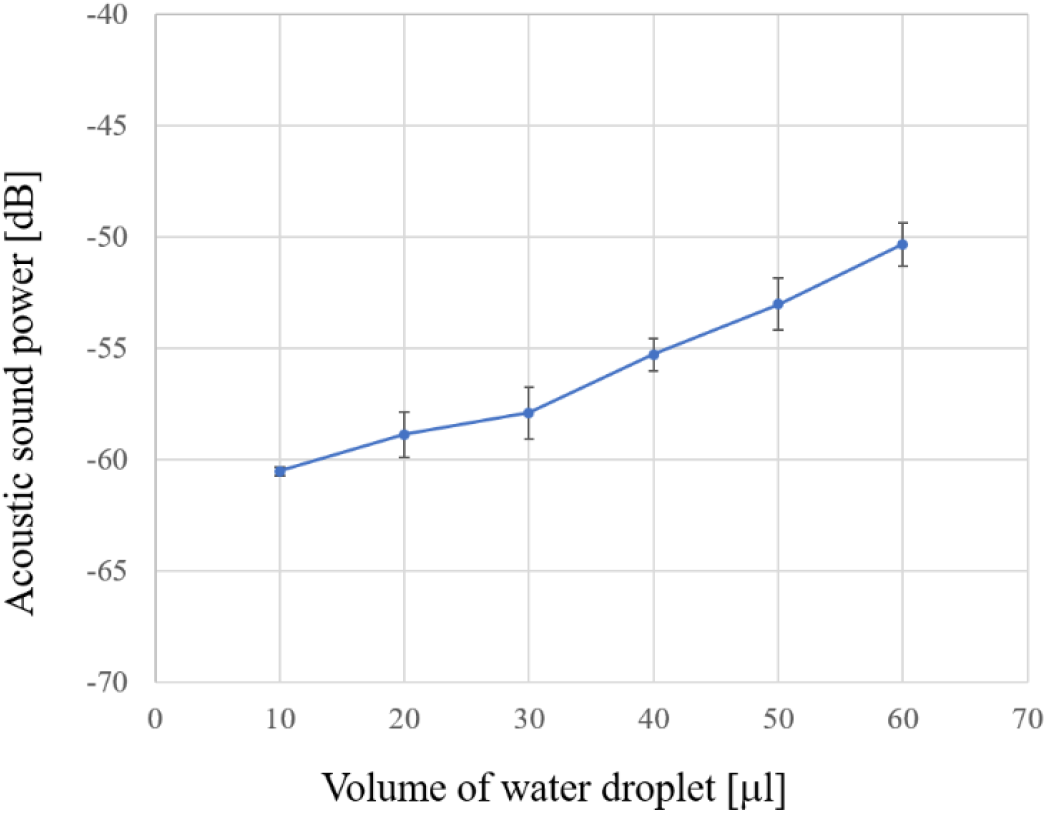
Dependence of averaged power within the 10 Hz to 5000 Hz range on the water droplet volume.

Next, possible effect of the guanine platelets from fish scale was investigated, as shown in Fig. 5. 40 μl of water droplet with and without the guanine platelets was placed on the membrane, and the power spectrum of the transformed sound was recorded. Compared to the distilled water droplet, the biogenic guanine platelet floating in water had an effect of enhancing the sound in the range of 10 Hz to 600 Hz, i.e. bass sound.

**Fig. 5.**
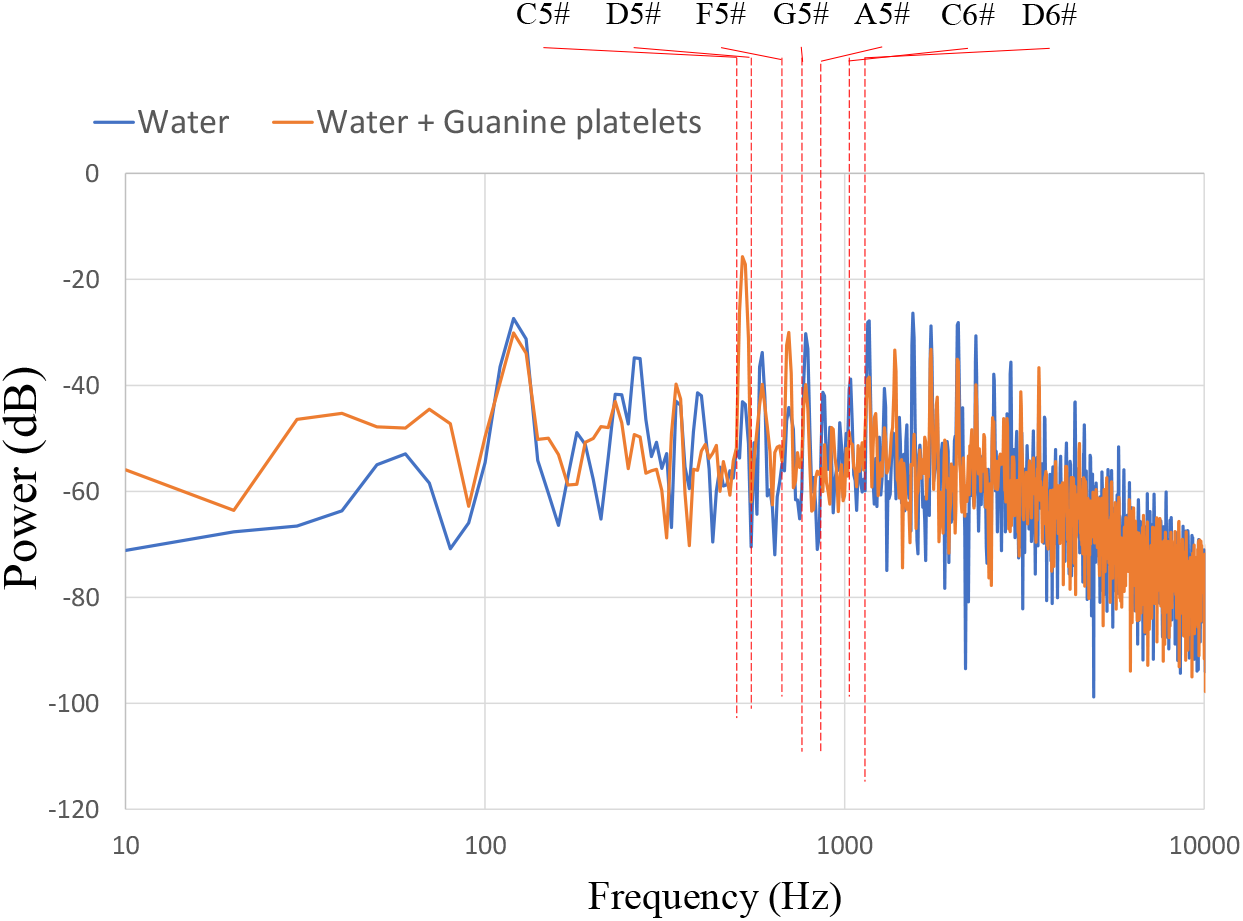
Effect of the biogenic guanine platelets floating in water on the power within the 10 Hz to 10000 Hz range on the water droplet volume.

Finally, we discuss the mechanism of the sound reconstruction phenomenon obtained via light reflection from the water droplet. Figure 6(a) shows photographs of the edge of the water droplet with and without sound exposure. The droplet edge became obscured when the piezoelectric membrane sounded, as illustrated in the right panel of Fig. 6(a). We failed to hear the reconstructed sound via the light reflection mechanism when the water droplet was absent (Fig. 2(c)). Therefore, the water droplet played a primary role in reflecting light that contained sufficient information to enable the sound reconstruction. The edge of the water droplet should contribute to this signal transduction process. In addition, the entire droplet region may contribute to generation of the light when involving the sound signal.

**Fig. 6.**
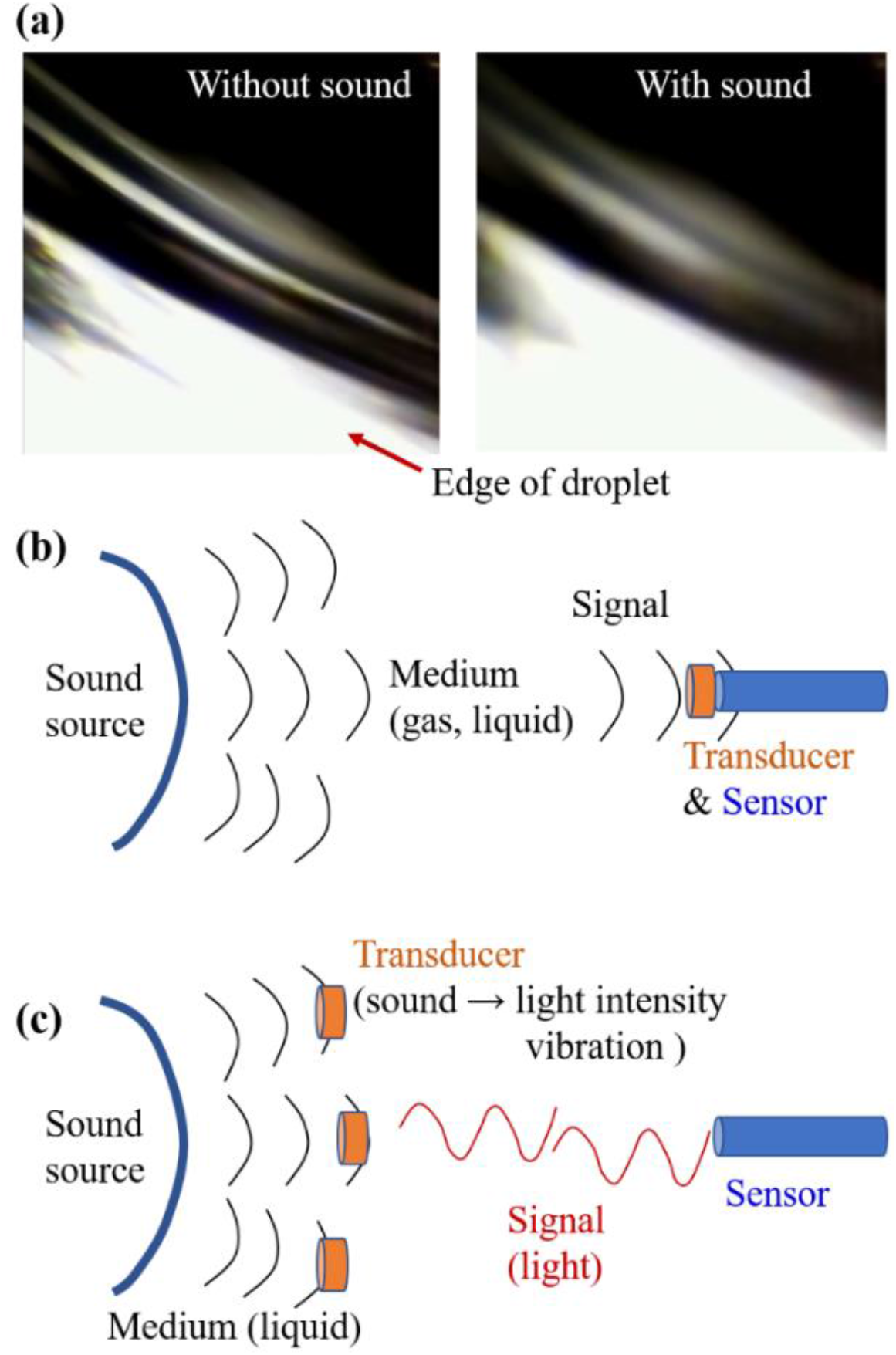
Water droplet acting as a transducer to transform frequency information from audible sound into light intensity. (a) Photographs of the edge of the water droplet with and without sound exposure. (b) Optical microphone transforming the sound into an optical signal at the edge of the sensor. (c) Principle of optical speaker, where the vibration of the water droplet transforms the sound information into light intensity and the frequency of this intensity.

Figure 6(b) and (c) illustrate the characteristics of the optical microphone strategy. Figure 6(b) illustrates the general principle of an optical microphone transforming sound into an optical signal at the edge of the sensor, e.g., an optical fiber sensor. The laser microphones used a laser for their light source and detected sound based on light interference. In that case, density fluctuations in the gas or the particles floating in the air can act as the transducer for the sound-to-light-signal transformation process.

This study proposes a method for sound detection using a small droplet of water which behaves like a speaker reflecting light carrying sounds. As illustrated in Fig. 6(c), the principle of this method was the vibration of the water droplet, which transformed sound information into light intensity and the frequency of this intensity. A liquid droplet adhering to the free surface of a vibrating object is useful for remote sensing of the audio signals involved in terms of the light intensity. Floating guanine platelets in the water had an effect of sound spectrum modulation, provably due to changing the vibration speed of the droplet. Because the guanine platelets have a high reflectivity and mechanically rotate in the water, a resonance frequency of the rotation and the applied sound can provide a specific power spectrum pattern.

## Conclusions

In conclusion, when a photosensor (PMT) was used, the analog output for the detected light produced a clear sound signal when the PMT collected the light reflected from the water droplet placed on the piezo-speaker that generated the sound. The acoustic features of the sound were modified by varying the volume of the water droplet. The power of the spectrum depended linearly on the volume of the water droplet.

The vibrating water droplet had a high-level capability for transformation of the acoustic features of the sound, comprising a chord composed of notes sounded by a musical instrument, into a light intensity that varied with frequencies ranging from 10 Hz to 5000 Hz. Adding guanine platelets obtained from goldfish scale modified the reflected light intensity at specific frequencies of acoustic wave. Consequently, the guanine platelets exhibited a micro mechanical actuation of the light-intensity-wave that carried the information of the audible sound frequency.

## Supporting information

SM_optical_speaker

## Supplementary materials

Description of the sounds referred in main text, Supplementary_Sound_1 ~ 3.

## Acknowledgment

This work was supported by JST-CREST “Advanced core technology for creation and practical utilization of innovative properties and functions based upon optics and photonic (Grant No.: JPMJCR16N1).” The author thanks David MacDonald, MSc, from Edanz (https://jp.edanz.com/ac) for editing a draft of this manuscript.

## Authors’ contributions

M. I. performed the experimental design, experiments, measurements and analyses. All parts of the manuscript and illustrations were prepared by M. I.

## References

1) J. Eargle, The Microphone Book-Second Edition, Focal Press (MA, U.S.A.) (2004)

2) G. M. Sessler, J. E. West, Foil Electret Microphone, J. Acoust. Soc. Am., 40, 6, 1433–1440 (1966)

3) G. S. K. Wong, T. E. W. Embleton, AIP Handbook of Condenser Microphone, AIP Press (1995)

4) E. Radcliffe, A. Naguib, W. M. Humphreys Jr, A novel design of a feedback-controlled optical microphone for aeroacoustics research, Meas. Sci. Technol., 21, 105208 (2010) https://doi.org/10.1088/0957-0233/21/10/105208

5) H. Takei, T. Hasegawa, K. Nakamura, S. Ueha, Measurement of intense ultrasound field in air using fiber optic probe, Jpn. J. Appl. Phys., 46, 4555 (2007) https://doi.org/10.1143/JJAP.46.4555

6) M. J. Murray, A. Davis, B. Redding, Fiber-wrapped mandrel microphone for low-noise acoustic measurements, J. Lightwave Tech., 36(16) 3205–3210 (2018)

7) J. P. F. Wooler, R. I. Crickmore, Fiber-optic microphones for battlefield acoustics, Appl. Opt. 46(13) 2486–2491 (2007)

8) K. Nakamura, S. Toda, M. Yamanouchi, A two-dimensional optical fibre microphone array with matrix-style data readout, Meas. Sci. Technol. 12 859 (2001) https://doi.org/10.1088/0957-0233/12/7/319

9) J. G. Choi, G. J. Diebold, Laser schlieren microphone for optoacoustic spectroscopy, Appl. Opt., 21(22) 4087–4091 (1982)

10) Y. Sonoda and M. Akazaki, Measurement of low-frequency ultrasonic waves by Fraunhofer diffraction, Jpn. J. Appl. Phys., 33, 3110–3114 (1994)

11) J. M. Moses, K. P. Trout, A simple laser microphone for classroom demonstration, The Phys. Teacher, 44 (9), 600 (2006)

12) B. Fisher, Optical microphone hears ultrasound, Nature Photonics, 10, 356–358 (2016)

13) C. Cai, K. Iwai, T. Nishiura, Y. Yamashita, Speech Enhancement for Optical Laser Microphone With Deep Neural Network, 2020 Asia-Pacific Signal and Information Processing Association Annual Summit and Conference (APSIPA ASC), 449–454 (2020)

14) A. Davis, M. Rubinstein, N. Wadhwa, G. J. Mysore, F. Durand, W. T. Freeman, The visual microphone: passive recovery of sound from video, ACM Transactions on Graphics, 33 (4), 79 (2014)

15) L. Zhou, J. He, W. Li, P. He, Q. Ye, B. Fu, P. Tao, C. Song, J. Wu, T. Deng, W. Shang, Butterfly wing hears sound: Acoustic detection using biophotonic nanostructure, Nano Letters, 19(4), 2627–2633, 2019

