## Supplementary material for "Modulation of optical speaker using biogenic guanine platelets floating in water": SM_optical_speaker

M. Iwasaka*

Hiroshima University; Kagamiyama 1-4-2 Higashihiroshima,
Hiroshima 739-8527, Japan.

Description of the sounds referred in main text:

**Supplementary_Sound_1**
Mp3 format
The sound (chord shown in Fig. 1(c)) is the original sound utilized in the experiments. The sound was pre-recorded using a PCM recorder.

**Supplementary_Sound_2**
Mp3 format
Supplementary data for Fig. 2. The sound was acquired via the water droplet on piezoelectric membrane under the exposure to side light illumination. Its power spectrum is shown in Fig. 2 (b). The time course of amplitude of the sound is shown in Fig. S1 where the piezoelectric membrane sounded in the middle phase ('Original sound on'). 'Off' means the period without the sound from piezoelectric membrane. White noise arose from the LED light source.

**Supplementary_Sound_3**
Mp3 format
Supplementary data for Fig. 3. The power spectrum of this sound is shown in Fig. 3 (a)-(f) where the volume of the water droplet was increased by adding every 10 μl of water using pipetting.

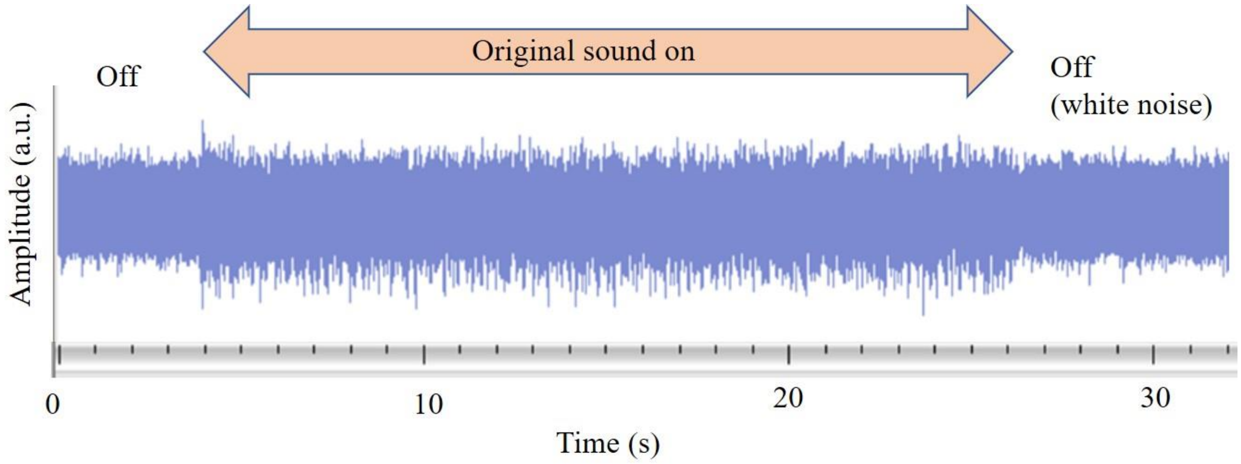


**Fig. S1.**

Time course of amplitude of the sound acquired via the water droplet on piezoelectric membrane under the exposure to side light illumination.

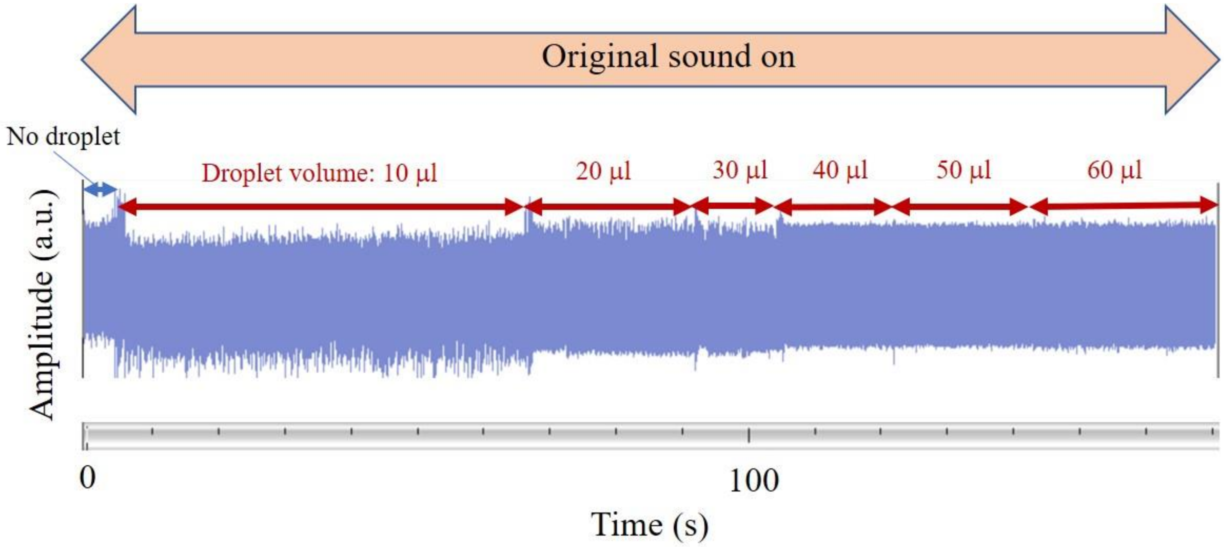


**Fig. S2.**

Time course of amplitude of the sound acquired via a water droplet when the volume of the droplet increased intermittently.
